# Comprehensive mapping of the 5′ and 3′ untranslated regions of *Aspergillus fumigatus* reveals new insights into gene regulation

**DOI:** 10.1101/2025.07.04.662582

**Authors:** Lukas Schrettenbrunner, Corinne Maufrais, Guilhem Janbon, Edward W. J. Wallace, Matthew G. Blango

**Affiliations:** Junior Research Group RNA Biology of Fungal Infections, Leibniz Institute for Natural Product Research and Infection Biology—Hans Knöll Institute (Leibniz-HKI), 07745 Jena, Germany; Institut Pasteur, Université Paris Cité, Unité Biologie des ARN des Pathogènes Fongiques, Département de Mycologie, F-75015, Paris, France; Institut Pasteur, Université Paris Cité, HUB Bioinformatique et Biostatistique, C3BI, USR 3756 IP CNRS, F-75015, Paris, France; University of Edinburgh, Institute for Cell Biology and Centre for Engineering Biology, School of Biological Sciences, University of Edinburgh, C.H. Waddington Building, Max Born Crescent, Edinburgh, Scotland, EH9 3BF, UK

## Abstract

In the twenty years since the first genome sequencing of *Aspergillus fumigatus*, the field has seen an explosion in both the number of sequenced genomes and our molecular understanding of this ubiquitous human fungal pathogen. Despite an improved knowledge of the *A. fumigatus* genome, we still know little about the transcriptome, with key regulatory sequences like the untranslated regions of mRNA based only on *in silico* predictions and bulk-RNA-seq. Here, we provide an improved description of 5′ and 3′ untranslated regions of *A. fumigatus* poly(A)-enriched RNA through experimental mapping of transcription start sites and polyadenylation sites using 5′ and 3′ End-Seq. We assigned high-quality 5′ ends to 2,747 genes (average length 126 nt), 3′ ends to 7,079 genes (average length 268 nt), and improved our understanding of the regulatory landscape of *A. fumigatus* gene expression. We leveraged the refined 5′ UTRs to identify upstream open reading frames and binding sites for important RNA binding proteins like the translational regulator Ssd1 and the 3′ UTRs to define binding sites for PUF proteins known to contribute to mRNA localization and regulation. Although a single isoform typically dominated expression, we observed 148 instances of alternative start sites and 1,675 alternative stop sites. Interestingly, we detected multiple examples of premature transcriptional termination, including the first evidence for promoter-proximal premature transcriptional termination in a member of the Eurotiomycetes. Ultimately, we provide a resource to the *Aspergillus* community and an accurate starting point for unravelling the complexities of gene regulation in an important human pathogen.

## INTRODUCTION

Transcriptome annotations are currently relied upon for myriad molecular investigations, yet in most cases these annotations themselves remain at best a work in progress. The value of a proper transcriptome annotation is obvious, as accuracy in transcription start and end sites facilitates definition of the regulatory elements responsible for the resultant RNA transcript and informs on transcription initiation, elongation, and termination. With correct transcript boundaries comes an improved understanding of upstream regulatory elements like promoters and enhancers; post-transcriptional regulatory elements like upstream open reading frames (uORFs) that ultimately contribute to mRNA stability and translation efficiency; and the coding potential of a given RNA (Aspden et al. 2023). Despite the clear advantages of an accurately annotated transcriptome and coding sequence, these resources are largely lacking from many eukaryotic pathogens and non-model organisms, a complicating factor we will partially remedy here for one noteworthy human fungal pathogen, *Aspergillus fumigatus*.

*A. fumigatus* is a World Health Organization critical priority fungal pathogen of the eurotiomycete class, highly diverged from the ascomycete model yeasts *Saccharomyces cerevisiae* and *Schizosaccharomyces pombe* (Li et al. 2021). *A. fumigatus* was first sequenced in 2005 using the reference strain and clinical isolate Af293 collected by postmortem lung biopsy of a neutropenic patient (Nierman et al. 2005). A second commonly used laboratory strain of the CEA10 lineage (A1163) was sequenced in 2008 (Fedorova et al. 2008), and the genome has been further improved and expanded by additional sequencing efforts, including a major expansion in sequenced strains through study of the pangenome (Barber et al. 2021; Bertuzzi et al. 2021; Horta et al. 2022; Lofgren et al. 2022; Rhodes et al. 2022).The first telomere-to-telomere assembly of *A. fumigatus* was published in 2022 using A1160 (a descendent strain of CEA10) and further broadened our understanding of the genomic architecture (Bowyer et al. 2022). The transcriptome of the *A. fumigatus* reference strain Af293 was initially annotated with the help of RNA-seq data and computational predictions, yielding better predicted UTRs than the more pathogenic CEA10 strain. To date, limited experimental evidence exists confirming the accuracy of the gene predictions for any of the *A. fumigatus* strains mentioned (Krappmann et al. 2004; Jöchl et al. 2008; Müller et al. 2012).

The boundaries of an RNA transcript, like a messenger RNA (mRNA) or long non-coding RNA (lncRNA), are bookended by a transcription start site at the 5′ end and a transcriptional terminator at the 3′ end. In mRNA and some lncRNA, a polyadenylation site (PAS) at the 3′ end of the transcript determines the site of cleavage prior to poly(A) addition. These features are not precisely defined, but instead are composed of multiple sites that provide regulatory versatility to the transcribed RNA. Much work has been done to assign transcript start sites (TSS) and PAS in a wide variety of organisms, particularly in model organisms like *S. cerevisiae* (McMillan et al. 2019; Dang et al. 2022). Software predictions are routinely performed with the support of RNA-seq data to assign untranslated regions (UTRs), but in many cases these predictions are incomplete and miss important regulatory features (Behr et al. 2013; Shenker et al. 2015).

Sequencing approaches designed to enrich for transcript ends provide a much more accurate picture of the mature RNA with UTRs than standard RNA-seq (Pelechano et al. 2014; Malabat et al. 2015; Afik et al. 2017). In many organisms, including fungi like *Cryptococcus neoformans* and *Neurospora crassa,* the selection of TSS dictates the inclusion or exclusion of regulatory elements such as mitochondrial localization signals or uORFs (Wallace et al. 2020). The resulting alternative protein forms have been termed echoforms to describe the case where a single gene encodes two or more differentially localized proteins (Yogev and Pines 2011; Bader et al. 2020). Recent work in *C. neoformans* provided a more complete characterization of TSSs and uncovered a novel transcription factor dictating alternative TSS selection (Dang et al. 2024). The best example of such regulation in *A. fumigatus* stems from studies of the conserved transcriptional activator <u>c</u>ross-<u>p</u>athway <u>c</u>ontrol <u>A</u> (CpcA; Gcn4p in yeast), which coordinates amino acid starvation responses (Krappmann et al. 2004). In *S. cerevisiae*, four uORFs in the 5′ leader of *GCN4* orchestrate translation (Gunišová and Valášek 2014), whereas the pathogenic yeast *Candida albicans* harbors a single regulatory uORF (Sundaram and Grant 2014). *A. fumigatus cpcA* has two uORFs but is also capable of autoregulation (Krappmann et al. 2004), highlighting the variability of such elements even at one vital, conserved locus.

Akin to transcription start sites, alternative polyadenylation (aPAS) sites are equally dynamic and important, with clear links to RNA stability (Tian and Manley 2017), RNA localization (Arora et al. 2022), and modulation of protein production (de Prisco et al. 2023), among others. In fungi, a range of roles have been described for aPAS, including regulation of stress resistance and virulence (Franceschetti et al. 2011; Graber et al. 2013). An under-estimated and often overlooked mechanism of gene regulation is premature transcription termination (PTT) (Kamieniarz-Gdula and Proudfoot 2019). PTT is most prevalent in metazoans as promoter proximal stalling of RNA polymerase II (Jonkers and Lis 2015). If stalling is not resolved, this can lead to PTT. Whether fungal specific PTT mechanism are functional in *A. fumigatus* is unknown. PTT can also be triggered by cryptic poly(A) signals often located within introns (Hoque et al. 2013). These examples demonstrate the regulatory potential of transcriptional start site and polyadenylation site selection.

Here, we improve our understanding of the *A. fumigatus* CEA10 transcriptome by performing 5′ and 3′ End-Seq coupled with RNA-seq to experimentally validate the ends of poly(A)-tailed RNA molecules. *A. fumigatus* has UTRs similar in length to ascomycetes like *S. pombe*, but relatively shorter 5′ UTRs and longer 3′UTRs compared to the basidiomycete pathogen *C. neoformans*. We uncover intriguing examples of premature transcription termination and other regulation that will serve as a resource to the community moving forward.

## RESULTS

### End-Seq defines 5′ and 3′ ends of *A. fumigatus* CEA10 transcripts

To determine baseline gene expression and transcript start (TSS) and end sites (TES) of *A. fumigatus* strain CEA10, 4 biological replicates of mycelium were grown in AMM liquid cultures for 24 h at 37°C (standard growth conditions) or 42°C (mild heat stress), followed by RNA-Seq and 5′ and 3′ End-Seq of the same RNA samples. DESeq2 analysis on the RNA-Seq data revealed minor changes due to heat stress, with 78 genes significantly downregulated and 75 upregulated (**Table S1**). No GO-terms were enriched for the downregulated genes; however, for the upregulated genes several enriched categories were identified using FungiDB.org (Amos et al. 2021). In the “Cellular Component” category four terms related to the cell wall were significantly upregulated (**Table S2**), consistent with the term “1,3-β-D-glucan metabolic process” that was observed in the “Biological Process” category. Collectively, these categories suggest minor cell wall stress, consistent with previous results from the literature indicating that *A. fumigatus* alters cell wall composition during heat stress (Fabri et al. 2021). Additionally, when we used FungiDB to search for enriched “Molecular Functions”, only the GO-term “HSP90 protein binding” was significantly enriched, hinting at a conserved heat shock response and adjustment of carbohydrate metabolism (Albrecht et al. 2010). As we only observed minor differences on the transcriptional level between the two growth conditions, we combined all replicates for the downstream End-Seq analysis to increase power.

5′ and 3′ End-Seq was performed on total RNA as described in **Fig. 1A**. Of note, for 5′ End-Seq, poly(A)-enriched RNA was heat-fragmented prior to phosphorylation and degradation of uncapped 5′ ends, whereas for 3′-End-Seq, total RNA was heat-fragmented prior to poly(A)-enrichment. 5′-End-Seq led to an enrichment upstream of many annotated start codons; however, there were also substantial reads mapping throughout the CDS and adjacent non-coding regions, whereas the resulting 3′-End-Seq reads were primarily located after the annotated end of the respective coding sequence (CDS), as expected.

**Figure 1:**
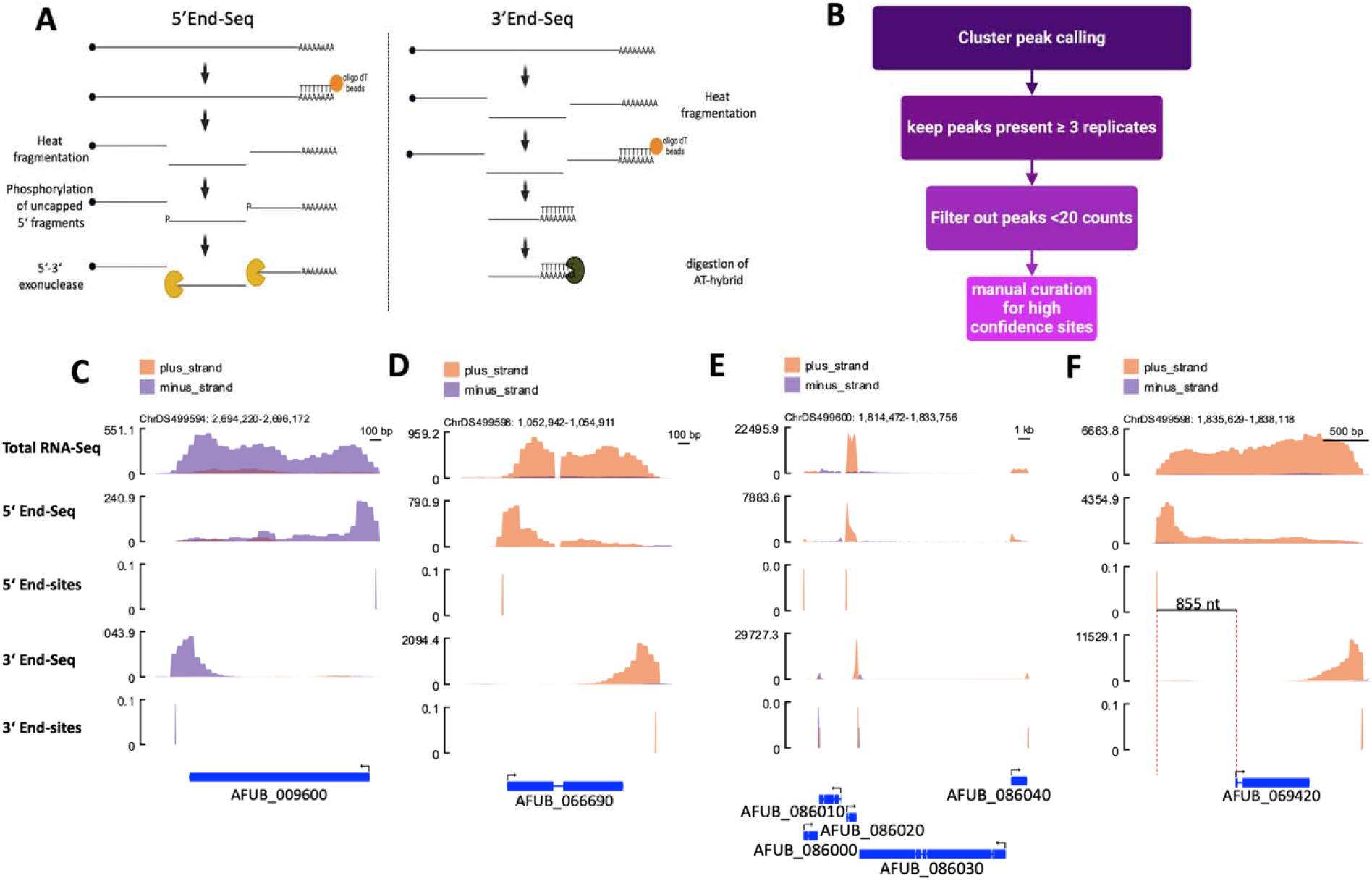
End-Seq defines UTRs in *A. fumigatus*. (A) Graphical description of the sequencing strategy using commercially available 5′ and 3′ End-Seq kits from Eclipse Bioinnovations (USA). For 5′ End-Seq we combined three biological replicates of fungus grown at 37°C and 42°C each for a total of six replicates, and for 3′ End-Seq we had four biological replicates for each condition resulting in eight total replicates. Created in BioRender. Blango, M. (2025) https://biorender.com/lguy6bt. (B) Schematic depiction of the filtering strategy to achieve high confidence sites. Created in BioRender. Blango, M. (2025) https://biorender.com/j5p9m7c. (C-F) Example plots of mRNA-Seq, 5′ End-Seq, filtered 5′ End-sites, 3′ End-Seq, and filtered 3′ End-sites. Orange indicates reads aligning to the + strand while purple indicates the – strand. The annotated AUG is indicated by an arrow, the CDS is indicated by the thick blue line, and introns by a thin blue line. The y-axis indicates total read-counts for “-Seq” panels; for “-sites” panels the y-axis is artificial as “sites” do not have a count. Read tracks were smoothened to improve viewability. Plots show (C) *AFUB_009600*, an mRNA splicing factor; (D) *AFUB_066690* encoding Aspf2, a major allergen of *A. fumigatus*; (E) genes of the pseurotin A biosynthetic gene cluster; and (F) *AFUB_069420* encoding CpcA with the length of the newly defined 5′ UTR indicated.

To determine the most prominent transcript end site, we extracted the 5′ and 3′ end positions of the respective reads and applied a previously established peak-calling algorithm (Dang et al. 2024). We kept sites identified in at least three replicates and a minimum of 20 total reads (**Fig. 1B**). The remaining sites were assigned to the closest gene on the same strand. All sites found within the annotated gene (0 nt distance from nearest annotated gene) or further than 800 nt from the annotated translational start or stop site were manually assessed for contiguous read coverage in the RNA-seq analysis indicative of a true TSS/TES. We then assigned each end site a confidence value, of “3”, “2”, or “1”, with “3” representing high-confidence sites, “2” indicating a probable end site requiring experimental validation, and “1” delineating an unlikely end site of an annotated gene in CEA10 (**Table S3**).

After manual curation, 2897 TSS and 9460 TES were assigned a high confidence value of “3”, covering 27.1% and 69.9% of 10,124 annotated genes in the CEA10 genome (ASM15014v1), respectively. Two representative examples of genes (e.g., *AFUB_009600*, a splicing factor; and *AFUB_066690*, encoding the major *A. fumigatus* allergen Aspf2 (Dasari et al. 2018)) with annotated TSS and TES sites are shown in **Fig. 1C-D**. For comparison, we performed a lift over of genome annotation coordinates from reference strain Af293 to CEA10 using the flo software package (https://github.com/wurmlab/flo), which generally improved the annotation; however, we also observed cases where the predicted UTRs do not match the experimental validation. In **Fig. 1E** the annotated start and stop sites of the pseurotin A biosynthetic gene cluster are shown, indicating the ability of End-Seq analysis to provide useful context even in gene dense loci. As an additional positive control, we confirmed the two uORF-containing 5′ end of the *cpcA* gene of *A. fumigatus* to occur at position −855, very similar to previous reports of a TSS at position −858 (**Fig. 1F, Table S3**; (Krappmann et al. 2004)). Building on this analysis, we extracted the sequences from the high-confidence 5′ UTRs and scanned for upstream start codons (uAUGs). 1180/2897 genes contained at least one uAUG (**Table S4**), suggesting vast, largely uninvestigated regulatory potential.

### A single mRNA isoform typically dominates *A. fumigatus* gene expression

The 2897 TSS and 9460 TES were annotated to 2747 and 7079 unique genes, respectively, resulting in 148 genes with alternative 5′ UTRs and 1675 with alternative 3′ UTRs based on the high-confidence ends. 2599 genes had only one TSS and 5404 genes only one TES. This suggests that the majority of *A. fumigatus* genes are transcribed as a single isoform without alternative TES and TSS under standard growth conditions. Globally, 3′ UTR variants were more common (∼25%) than 5′ UTR variants (∼5%). Among the genes exhibiting alternative TSS/TES, most have only one additional alternative (145 genes with two alternative 5′ UTRs; 1354 genes with two alternative 3′ UTRs) (**Fig. 2A-C**). A few genes have several potential UTR isoforms, for example 3 genes exhibit 3 alternative 5′ UTRs; 293 genes had 3 alternative 3′ UTRs, 87 with 4, 30 with 5, 9 with 6, 1 each with 7 and 9 3′ UTRs (see example in **Fig. 2D**). When there was an alternative TSS/TES, this usually correlated with an increase/drop in transcript level with respect to the main start or stop site, which can be observed in the “totalRNA” track (**Fig. 2**).

**Figure 2:**
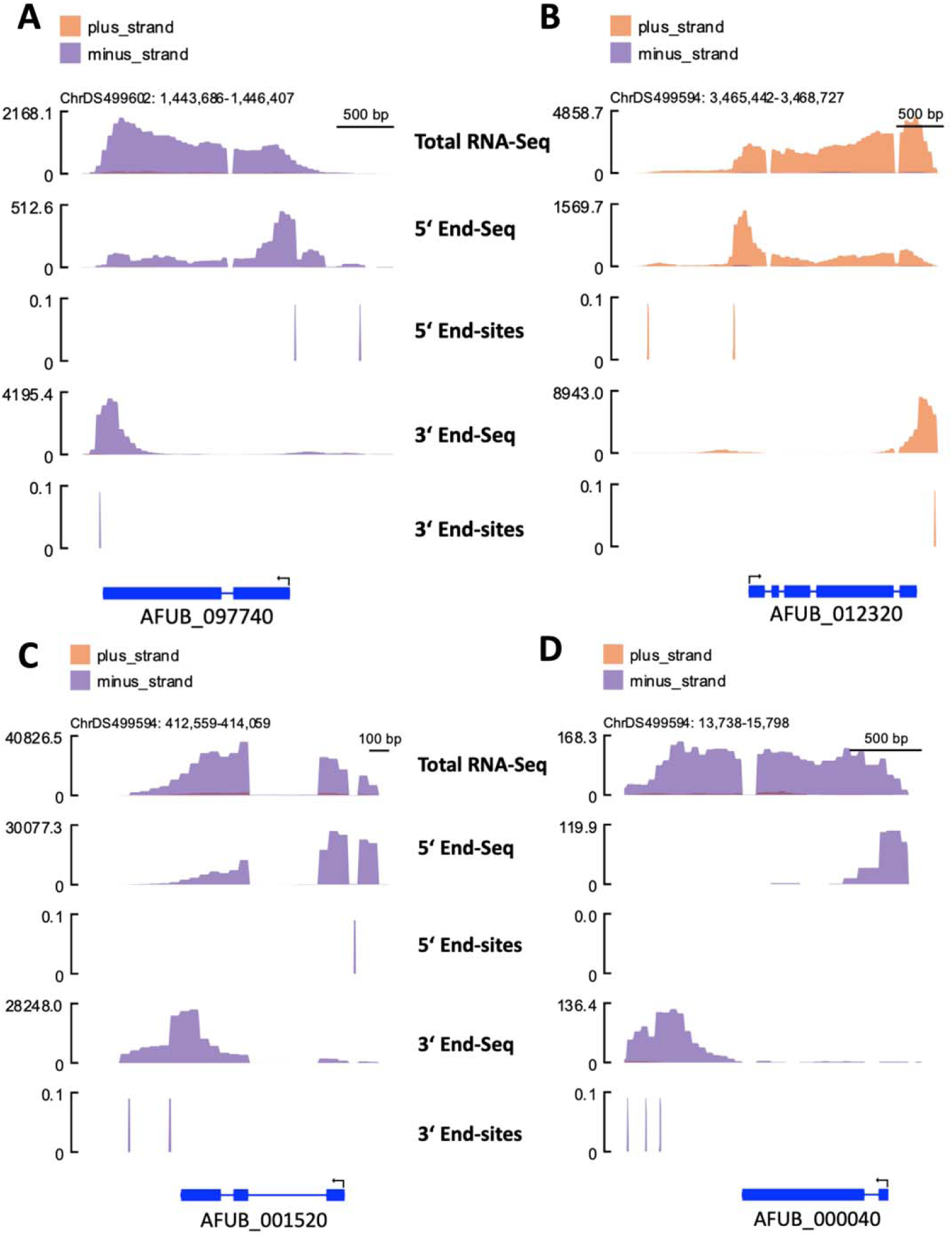
Multiple genes have transcripts with observed alternative 5_′_ and 3_′_ RNA end sites. (A) Read alignment of *AFUB_097740* gene encoding a protein of unknown function, (B) *AFUB_012320* encoding a putative nitrate transporter, (C) *AFUB_001520* encoding 60S ribosomal protein L31e, and (D) *AFUB_000040* encoding protein with predicted RNA polymerase II transcription factor activity. Read alignment of several genes; the y-axis indicates total read-counts for “-Seq” panels; for “-sites” panels the y-axis is artificial as “sites” do not have a count. Read tracks were smoothened to improve viewability.

### *A. fumigatus* has longer 3′ UTRs than other reported fungi, despite comparable 5_′_UTRs

We analyzed high-confidence end sites (value of 3) to determine the average length of the UTRs in *A. fumigatus* CEA10 (**Fig. 3A; Table S3**). The mean length of 5′ UTRs was 126 nt with a median length of 49 nt, whereas the mean length of the 3′ UTRs was slightly longer at 268 nt, with a median length of 184 nt. Compared to UTR lengths in *C. neoformans* and *S. pombe*, *A. fumigatus* 5′ UTRs were slightly shorter (177 nt in *C. neoformans* and 152 nt in *S. pombe* vs 126 nt mean length in *A. fumigatus*) but longer than *C. albicans* 5′ UTRs, which were 88 nt long on average. This may be partially attributed to the large number of “internal start-sites” that bias the 5′ UTR to be shorter in *A. fumigatus*. The *A. fumigatus* 3′ UTRs were notably longer than those of *C. neoformans, S. cerevisiae, S. pombe,* and *C, albicans* (186, 144, 169, and 84 nt vs 268 nt mean length; **Fig. 3B**). We performed an additional control analysis excluding any UTRs with a calculated length of zero relative to the annotated gene and determined 5′ UTRs to be 218 nt (mean) with a median length of 135 nt and 3′ UTRs to be 282 nt (mean) with a median of 195 nt.

**Figure 3:**
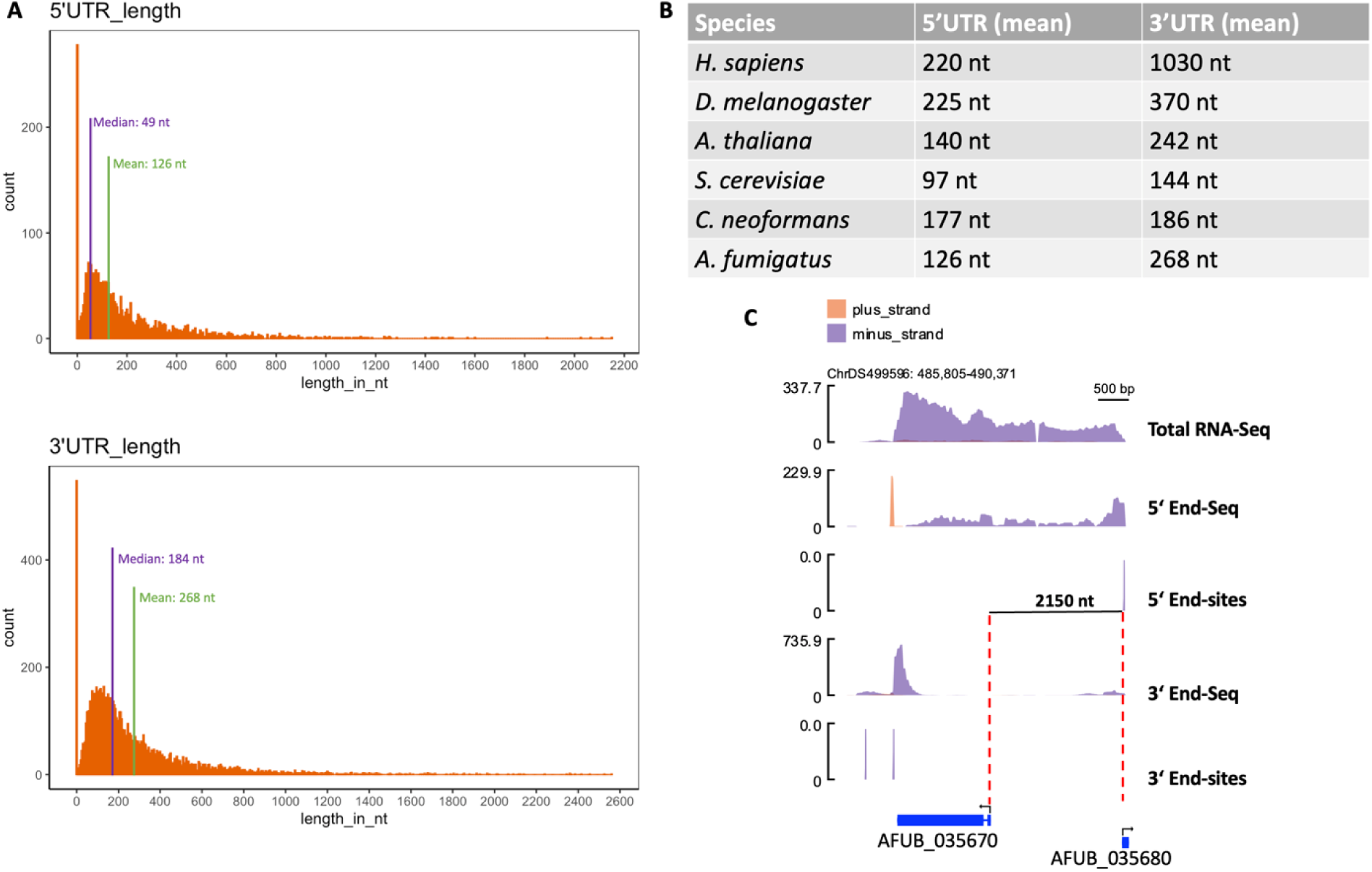
*A. fumigatus* exhibits 5_′_ UTRs of typical length, but relatively longer 3_′_ UTRs than other fungi. (A) Histograms (bin size equals 5) of UTR length distribution based on the observed high-confidence 5′ and 3′ poly(A)-enriched RNA ends. (B) Comparison of average UTR lengths with other eukaryotes (Graber et al. 1999; Misra et al. 2002; Chen et al. 2011; Srivastava et al. 2018; Wallace et al. 2020; Hong and Jeong 2023). (C) Transcriptomic snapshot of *AFUB_035670* together with the measured 5′ and 3′ End sites; the 5′ UTR of *AFUB_035670* was determined to be 2150 nt long compared to the prior prediction of 703 nt in Af293. Read tracks were smoothened to improve viewability.

We observed a number of TSS (and to a much lesser extent TES) within the boundaries of the annotated CDS of genes, likely due to incorrect annotation. In most cases the reported TSS was confirmed by total RNA-seq data and exhibited transcription downstream of the annotated start codon. In these cases, the current CDS annotation prediction is likely too long and includes an upstream (in-frame) start-codon (for an example see **Fig. S1**), which could be utilized under different growth conditions. We often observed a few reads aligning to the upstream region, indicating that there could be an alternative TSS (as in **Fig. 2B**) although these events typically did not meet our threshold for inclusion as *bona fide* TSS. The inclusion of an upstream start codon has previously been linked to signal peptides that alter protein localization (Natsoulis et al. 1986; Mireau et al. 1996; Mudge et al. 1998; Wallace et al. 2020). One particularly interesting example warranting future study is the 2150-nt-long 5′ UTR of *AFUB_035670*, which contains both a UTR intron and 30 uAUGs, and coincides with the promoter of an antisense neighboring gene, *AFUB_035680* (**Fig. 3C**). In silico prediction of the coding sequence using the longer 5′ UTR did not reveal an obvious contiguous open reading frame, suggestive of a regulatory role rather than misannotation in this case. Similarly, the *AFUB_067240* gene showed the highest number of uAUGs at 34 occurences in its 2111-nt 5′ UTR. In fact, 275/1180 had five or more uAUGS (**Table S4**), which makes it unlikely that the annotated start codon is used and suggests a misannotated start codon or unusual regulation. Further work including ribosome profiling will be needed to understand these unusual examples.

### Shortest and longest UTRs reveal novel features of *A. fumigatus* gene regulation

We divided the high-confidence UTRs according to their length and assessed the 10% shortest and longest regions (**Fig. S2**). A GO-term analysis of the 10% shortest 5′ UTRs showed a significant enrichment of ribosomal genes (e.g., GO-terms: “structural constituent of ribosome”, “ribosome”, “translation”) and genes associated with mitochondrial membranes (“mitochondrial respiratory chain complex III assembly”) (**Fig. S2A**; **Table S5**). These findings are consistent with the idea that genes requiring high levels of translation, like those encoding ribosomal proteins, harbor shorter UTRs as a means to optimize translational output (reviewed in (Petibon et al. 2021)). Among the genes with the 10% longest 5′ UTRs, phosphatases and kinases were especially common (e.g., “protein phosphorylation”, “protein kinase activity”, “Transferase activity, transferring phosphorus-containing groups”) (**Fig. S2B**). Genes involved RNA/DNA interaction were also enriched (e.g., “DNA-binding transcription factor activity”, “Nucleic acid binding”, “chromatin binding”). As longer UTRs are often postulated to possess additional regulatory capacity, these genes with long UTRs may be a particularly appealing place to find novel post-transcriptional regulatory mechanisms in *A. fumigatus* and related species.

We next extracted genomic sequences +/- 25 nt from the transcription start site of the 5′ UTRs and predicted motifs using STREME (version 5.5.7) (Bailey 2021). We observed several motifs with significant albeit limited enrichment (**Fig. S3A**). When we did the same STREME analysis on the 10% shortest and longest 5′ UTRs, 3 different motifs were identified in each case (**Fig. S3B,C**); however, the enrichment for these features was also minimal at best, indicating that a more sophisticated analysis will be required to tease apart these regulatory features of transcription in *A. fumigatus*. Next, WEB-logos were created from these sequences to assess potential sequence bias around the TSS. For both long and short UTRs, we found an enrichment of a TG at positions −1 and 0 (**Fig. S3D**). Interestingly this enrichment of TG was not observed when the sequences for all identified high confidence start sites were used. Instead, an enrichment for a CA at positions 1 and 2 was observed, suggesting some specificity in the longest and shortest UTRs (**Fig. S3D**). Of note, while A and G were slightly more prevalent upstream and downstream of the TSS in the short and long UTR genes, C and T were slightly more prevalent upstream and downstream of positions −1 and 0 when all TSS were considered. Interestingly, in *S. cerevisiae* T-richness upstream of the TSS is positively correlated with promoter activity (Lubliner et al. 2013).

No GO-term was enriched among the genes with the 10% shortest 3′ UTRs. Enriched GO-terms for the 10% longest 3′ UTRs could generally be grouped into 3 categories: 1) Transcription (e.g., “Regulation of DNA templated transcription”, “DNA-binding transcription factor activity, RNA polymerase II specific””, “Sequence specific DNA binding”); 2) phosphatases and kinases (e.g., “Protein kinase activity”, “Transferase activity, transferring phosphorus containing groups”, “Protein phosphorylation”); and 3) signaling (e.g., “G protein-coupled receptor signaling pathway”, “signal transduction”, “Heterotrimeric G-protein complex”)(**Fig. S2C**). Collectively, assessment of the longest and shortest UTRs revealed a reliance on short UTRs for ribosomal genes and longer UTRs for genes kinases and phosphatases.

### Known RNA binding motifs are enriched in UTRs of expected client genes

Ssd1/SsdA/gul-1/Sts5 is a fungal specific RNA-binding protein that regulates growth and cell wall synthesis (Ballou et al. 2020; Hall and Wallace 2022). *A. fumigatus* SsdA is critical for regulation of morphology by the CotA kinase (Martin-Vicente et al. 2024). The binding motif (CNYTCNYT) of *S. cerevisiae* Ssd1 is well characterized and highly enriched in the 5′ UTRs of many mRNAs (Hogan et al. 2008; Bayne et al. 2021). For efficient binding the motif should occur at least twice within the 5′ UTR (Bayne et al. 2021). We scanned our newly defined 5′ UTR sequences for instances with ≥2 Ssd1 binding motifs and found 527 such genes (**Fig. 4A**). *AFUB_*063890 (encoding the ortholog of *S. cerevisiae* Ecm33) held the most 5′ UTR Ssd1-binding sites with 14. Ecm33 is a GPI-anchored cell wall protein and known client mRNA of Ssd1 in *S. cerevisiae* (Li et al. 2013). A GO-term analysis using the 527 genes with ≥2 Ssd1 binding sites and a background list of genes with high-confidence 5′ UTRs revealed an enrichment of the categories “fungal-type cell wall”, “fungal-type cell wall organization”, “carbohydrate metabolic process”, “nucleic acid binding” and “DNA binding” (**Fig. 4B**), consistent with the literature (Hogan et al. 2008; Bayne et al. 2021). We also confirmed that the Ssd1 binding motif was indeed enriched in 5′ UTRs by shuffling the 5′ UTR sequences and in parallel running the same analysis on 3′ UTR sequences. In both cases this showed a significantly reduced number of motifs (**Fig. 4C**).

**Figure 4:**
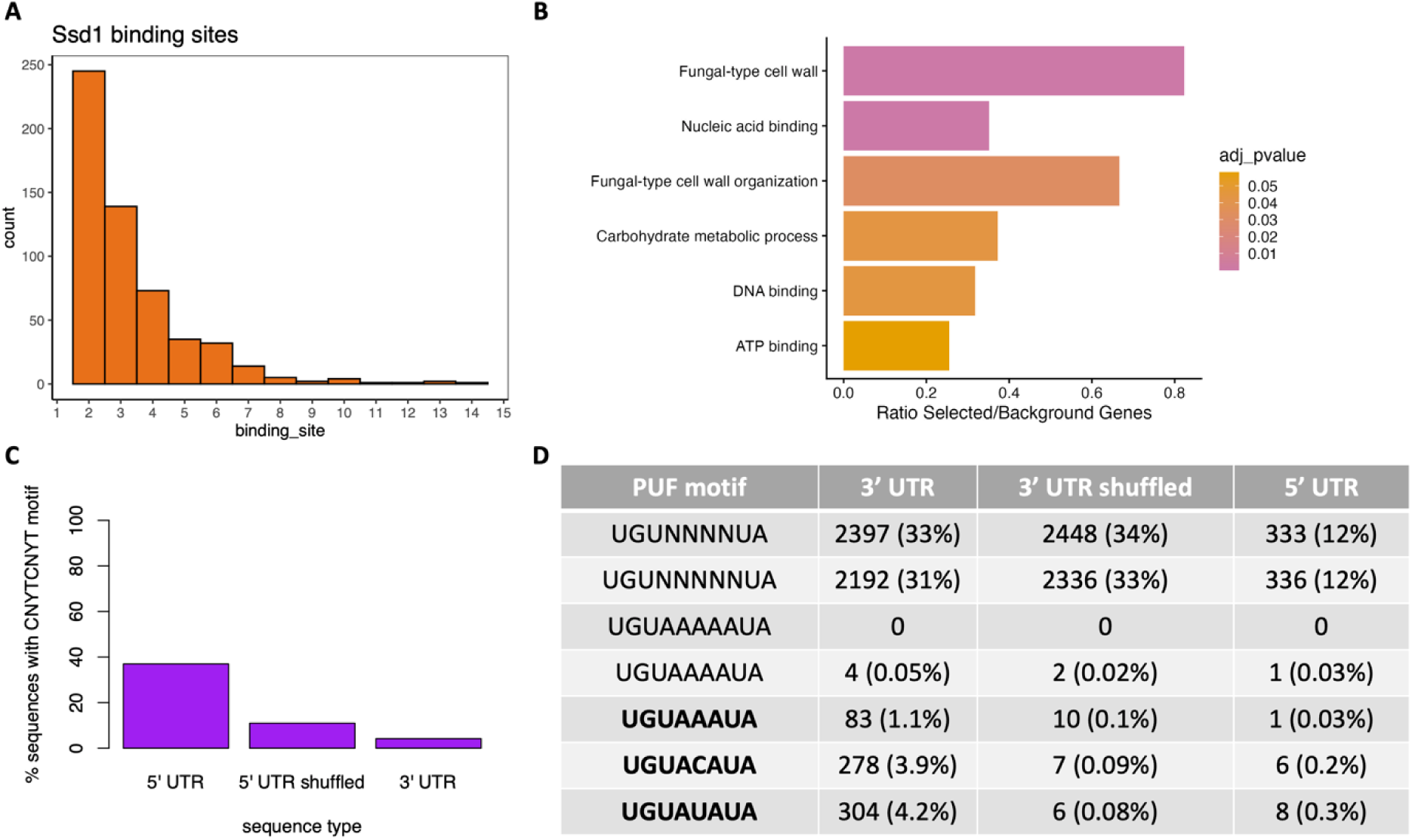
Ssd1 and PUF binding sites are enriched in 5′ and 3′ UTRs, respectively. A) Histogram showing the number of Ssd1 binding motif in newly defined 5′ UTRs. Only genes with at least 2 motif occurrences were plotted. (B) GO-term analysis of the same genes as in (A) with a supplied background list of all genes with an identified 5′ UTR in our study. GO-term analysis was done using https://fungifun3.hki-jena.de/. (C) percent of CNYTCNYT motif occurrence in different sequence types. For this graph a single occurrence was also counted. (D) Potential PUF protein family binding motif occurrence in different sequence types. Indicated are the number of sequences with the respective motif and the percent of the total number of input sequences in brackets. Sequence motifs were taken from (Hogan et al. 2015; Wilinski et al. 2017).

The Pumilio and FBF (PUF) family RNA binding proteins mainly interact with the 3′ UTRs of mRNAs encoding for membrane associated proteins (Wang et al. 2018; Murante and Hogan 2019). The literature suggests several potential binding motifs, and we scanned our identified 3′ UTR sequences for their occurrence (Hogan et al. 2008; Wilinski et al. 2017; Sadée et al. 2022). Only the motif UGUA[ACU]AUA resulted in enriched binding sites over the background (5′ UTR sequences and shuffled 3′ UTR sequences) (**Fig. 4D**). We identified 653 genes with at least one of the three sequences occurring in a 3′ UTR. A GO-term analysis with all genes yielded no enrichment; however, when only the 19 genes with 2 binding sites were considered (2 binding sites was the maximum observed) several GO terms were significantly enriched. The terms “uniporter activity”, “mitochondrial calcium ion homeostasis”, and “calcium channel activity” were comparable to binding patterns of PUF family proteins in other species (García-Rodríguez et al. 2007), but the small number of hits in this category may suggest further refinement is necessary to understand PUF protein biology in *A. fumigatus*.

Overall, the enrichment of known protein-binding motifs in newly annotated UTR sequences argues that these sequences can be used to discover new binding motifs and regulatory functions.

### End-Seq reveals numerous cases of putative premature (intronic) transcription termination

We believe that the dataset provided here will prove a tool for hypothesis generation moving forward. One compelling area for future study centers on the 484 (of 9460 total) identified TESs located within the predicted CDS of genes. These events can be classified as cases of 1) early termination shortly after the TSS, 2) slightly shorter mRNA isoforms near the predicted TES likely resulting from poor annotation, and 3) transcripts that are significantly shortened, potentially resulting in truncated peptides. We can also differentiate between intronic and exonic sites. While exonic transcription termination sites most likely lead to degradation of the transcript by mechanisms like non-stop decay, intronic termination can yield shortened proteins (van Hoof et al. 2002; Vasudevan et al. 2002; Kamieniarz-Gdula and Proudfoot 2019). Out of the 484 3′ sites located within CDS, 154 are located either within an intron or adjacent to an intron that is alternatively spliced.

A particularly interesting example comes from the *AFUB_001550* transcript. AFUB_001550 is the predicted ortholog of *S. cerevisiae* ENV9, an oxidoreductase that plays an important role in lipid droplet morphology (Siddiqah et al. 2017). The pre-mRNA consists of 4 exons and 3 introns, where the first two introns are short and the third is relatively long at 274 nucleotides, with an in-frame stop-codon. Our sequencing data suggest that transcription is primarily terminated within the last intron, and only a fraction of reads reach the purported 3′ UTR. This can be observed in the total-poly(A)-RNA sequencing track as a drop in reads after the last intron (**Fig. 5A,B**). In *S. cerevisiae*, ENV9 functions as a short-chain dehydrogenase and protein domain predictions for full-length AFUB_001550 reveal similar short-chain dehydrogenase domains. In both orthologs, these domains are in the C-terminal half of the protein and predicted to be absent in the truncated peptide produced from transcript using the TES in intron 3. In *S. cerevisiae*, ENV9 is important for lipid-droplet morphology when the yeast is grown on poor carbon sources like glycerol or ethanol, with the C-terminus playing an important role in membrane anchoring (Siddiqah et al. 2017). The short transcript produced by *A. fumigatus* may hint at an additional layer of regulation during transcription to limit membrane tethering or dehydrogenase activity of AFUB_001550 under favorable growth conditions or alternatively suggest that the gene is undergoing negative (purifying) selection.

**Figure 5:**
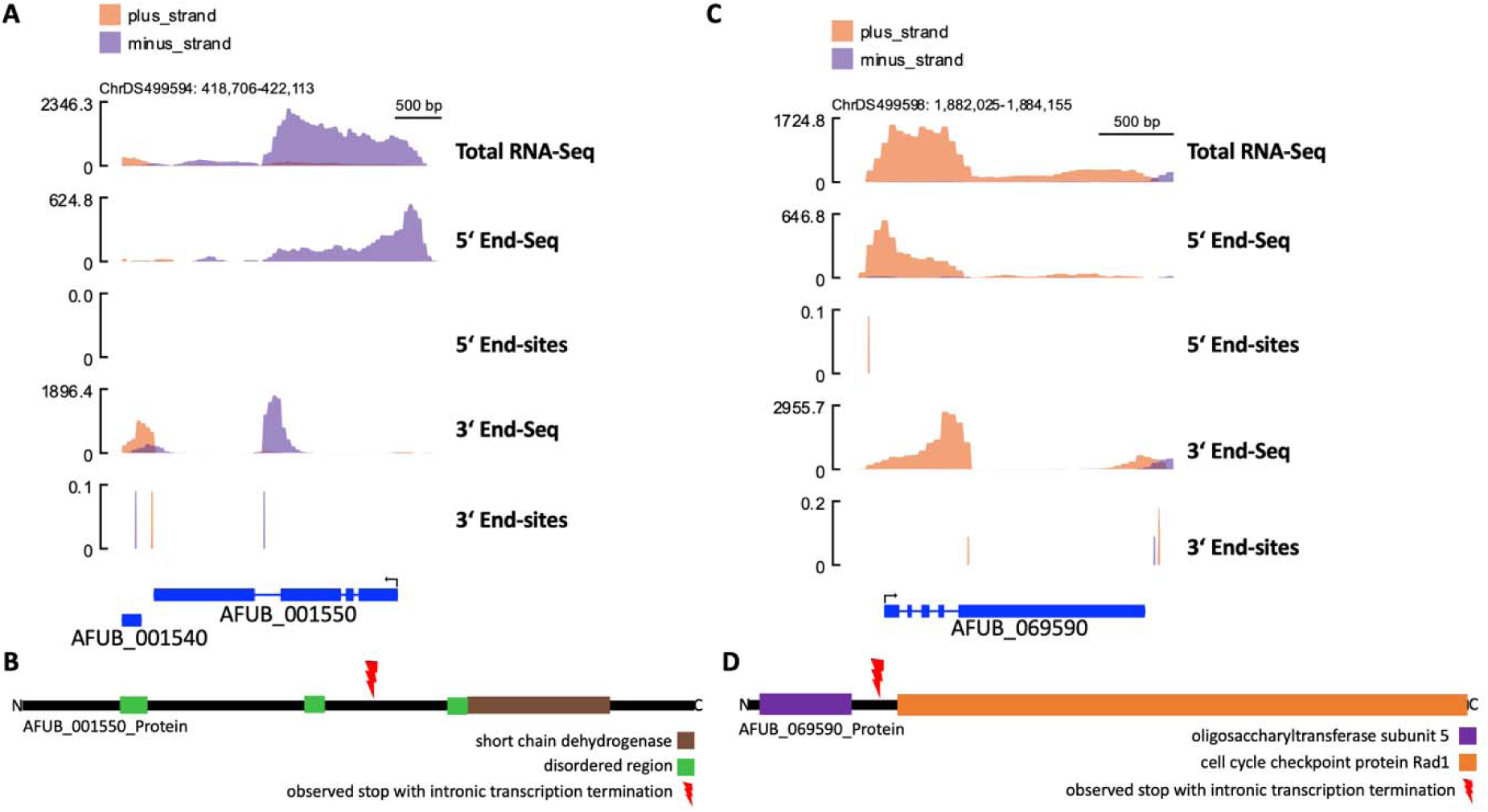
3′-End-seq reveals examples of premature termination in *A. fumigatus*. (A) Read alignment of *AFUB_001550* encoding ortholog of *S. cerevisiae* ENV9; the y-axis indicates total read-counts for “-Seq” panels; for “-sites” panels the y-axis is artificial as “sites” do not have a count. (B) Schematic depiction of the predicted protein domain of AFUB_001550 using data from InterPro; the location of the observed internal stop-site is indicated. (C) Read alignment of the previously uncharacterized *AFUB_069590* gene; the y-axis indicates total read-counts for “-Seq” panels; for “-sites” panels the y-axis is artificial as “sites” do not have a count. (D) Schematic depiction of the predicted protein domain of AFUB_069590 using data from InterPro; the approximate location of the observed internal stop-site is indicated. Read tracks were smoothened to improve viewability.

A second similar example is the uncharacterized gene *AFUB_069590*, which consists of 5 exons and 4 introns (**Fig. 5C**). Most transcripts in our analysis ended in the last intron, which appears to be mostly retained. Protein domain prediction identifies a Rad1 domain, typical of cell cycle control proteins (Marathi et al. 1998), in the last exon that would be lost using the premature termination site (**Fig. 5D**). The first 4 (relatively short) exons encode a predicted oligosaccharyltransferase domain; however, as in the example for *AFUB_001550* above, the exact function of these domains in tandem requires further study.

### *A. fumigatus* exhibits putative promoter proximal transcription termination

As premature transcription termination is a relevant regulatory mechanism across multiple kingdoms (Kamieniarz-Gdula and Proudfoot 2019), we next assessed putative PTT sites in our dataset. We identified 42 genes with putative promoter proximal PTT using our high-confidence end sites, but it is noteworthy that there are certainly more potential promoter proximal PTT sites in the lower confidence groups, as these are often rare events that would not be expected to meet all our thresholds for high-confidence end sites. Additional examples were identified further downstream in the gene body (**Table S2** and **Fig. 5**), but here we focus on the promoter proximal examples. Many of our identified cases are likely derived from cryptic intronic PAS. For example, *AFUB_004020* exhibits both a detectable main TES and a minor alternative TES downstream of the annotated stop codon (**Fig. 6A**). According to our sequencing data, *AFUB_004020* also has a TES within the first intron, producing a transcript of only around 300 nt in length. Another interesting example is *AFUB_039430*, where the internal TES is in the third intron, the longest of the transcript (**Fig. 6B**). Here not only do we see reads for the 3′ ends but also a drop in total transcription around the same location as the internal TES. Although both examples shown in **Fig. 6** are also located close to or within introns, they are functionally different from the examples in **Fig. 5**. In the latter case, the premature transcription termination leads to a decrease in transcription and a shortened mRNA, while in the former no substantial drop in transcription after the premature transcription is observed. These putatively short RNAs could either be targeted for degradation (although they were stable enough to sequence), act as non-coding RNA, or even be translated into micropeptides (Kamieniarz-Gdula and Proudfoot 2019) to influence the *A. fumigatus* transcriptome/translatome. Additional effort will be required to fully understand the breadth and importance of PTT in *A. fumigatus* gene regulation.

**Figure 6:**
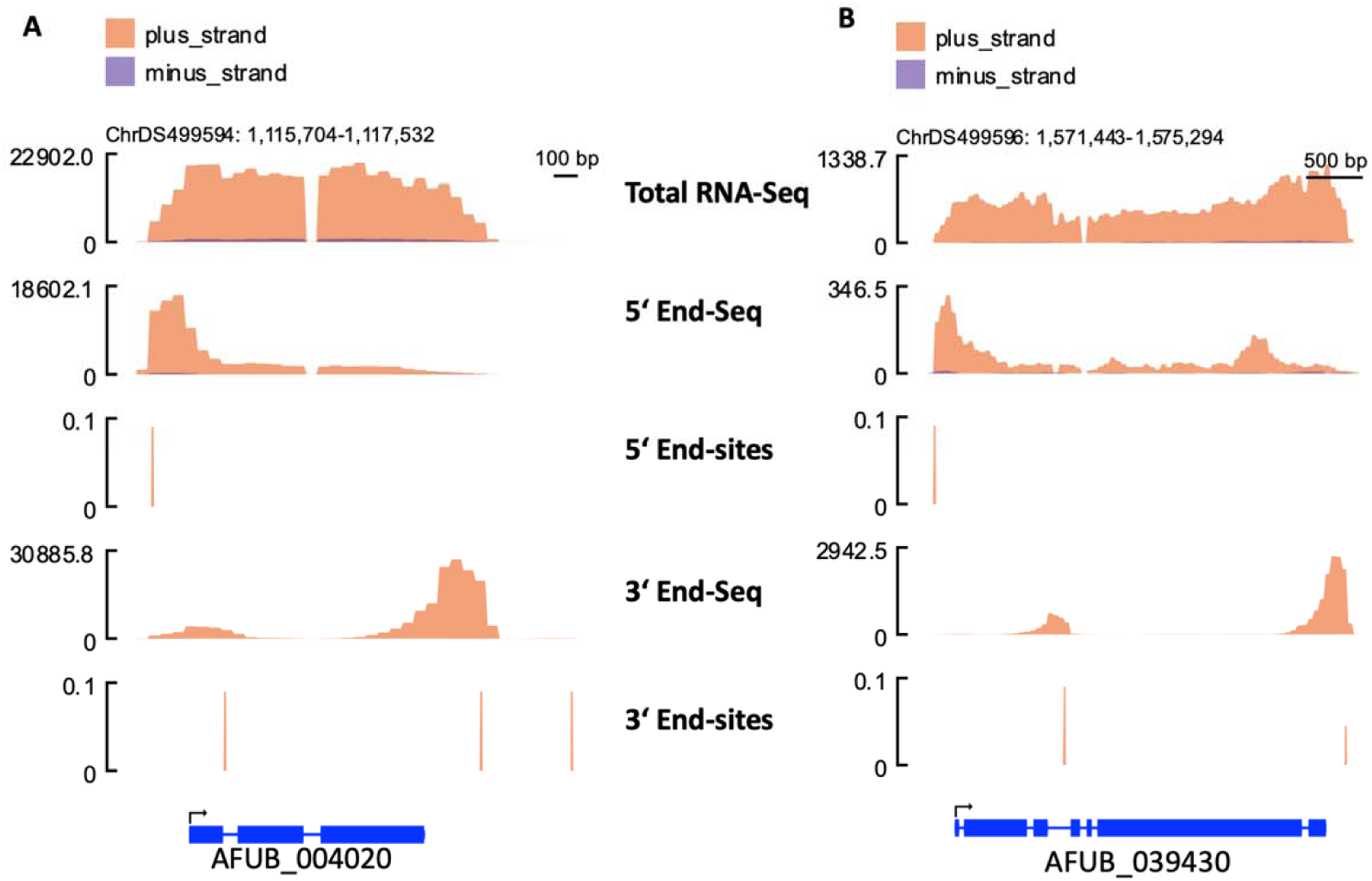
Promoter proximal transcription termination occurs in *A. fumigatus*. Read alignment of (A) *AFUB_004020* and (B) *AFUB_039430*, each encoding proteins of unknown function. The y-axis indicates total read-counts for “-Seq” panels; the y-axis is artificial for “-sites” panels as “sites” do not have a count. Read tracks were smoothened to improve viewability.

## DISCUSSION

The current public genome of the common laboratory strain CEA10 of *A. fumigatus* largely lacks experimental validation or even predictive information about the termini of RNA transcripts. Here, we provide an improved description of *A. fumigatus* CEA10 transcript start and end sites of poly(A)-tailed RNA from fungal mycelium. We were able to measure high-confidence TSS for 27.1% and TES for 69.9% of all annotated genes. The inclusion of middle- and potentially even low-confidence end sites in the supplemental materials (**Table S3**) increases these numbers further and will likely prove informative on a case-by-case basis for genes-of-interest.

We expect this resource to be useful to the community in several ways. First, a more complete map of the *A. fumigatus* UTRs will support improved molecular cloning projects in terms of creation of genetic deletion and overexpression strains. Second, a more accurate description of UTRs will facilitate our understanding of the regulatory potential of such elements, particularly as protein binding hubs. UTR-binding RNA binding proteins are well known to regulate the level of translation, in addition to the stability and localization of the RNA (Kurischko et al. 2011; Mendonsa et al. 2023).

The GO-term analysis performed on the genes with the shortest and longest UTRs confirmed that genes harboring a short 5′ UTR were enriched for ribosomal and mitochondrial genes, consistent with high translation. Genes with the longest UTRs on the other hand were enriched in GO-categories where additional regulation on the RNA level could be required, including kinases, phosphatases, metabolic genes, and transcription factors. Interestingly, the nucleotide frequency around the TSS of the long and short UTRs was quite different from the nucleotide frequency around all TSSs (**Fig. S3D**). There, we observed a higher frequency of Ts, which has been shown in *S. cerevisiae* to be positively correlated with promoter activity (Lubliner et al. 2013). This could suggest that the genes with short/long UTRs are under different regulatory pressures, although it is worth noting that there is a very enriched T at position −1 from the TSS.

While 3′-End-Seq reads mostly accumulated at the expected location close to the end of genes, 5′-End-Seq reads were typically found throughout the gene body. This is likely caused by the technical limitations of identifying 5′ ends using poly(A)-transcript enrichment, which requires selection of intact, full-length transcripts, followed by heat fragmentation, phosphorylation of non-5′-m^7^G-capped RNA, and finally enzymatic degradation of these phosphorylated fragments prior to library preparation (Afik et al. 2017). If the fragments are not fully phosphorylated and degraded, incorrect RNA fragments can be incorporated into the sequencing library. Nevertheless, for many genes we observed a clear enrichment of reads in proximity to the expected TSS. The nature of the poly(A)-enrichment and selection of TES meant that this dataset had lower noise, allowing for more accurate definition of end sites. With this dataset, we can more precisely define the *A. fumigatus* transcriptome, for example showing it to possess 5′ UTRs of 126-nt mean length similar to other fungi and slightly longer 3′ UTRs at 268 nt. These UTRs harbored numerous uAUGs and motifs for interaction with RNA binding proteins like SsdA and the PUF proteins. It will be interesting to see how UTR lengths change under stress or throughout *A. fumigatus* development in the future, perhaps supplemented by direct RNA sequencing (Depledge et al. 2019; Parker et al. 2020; Ibrahim et al. 2021) or ribosomal profiling approaches as performed previously (Wallace et al. 2020).

Previous RNA-end analyses have proven that transcription does not always perfectly start or stop (or undergo cleavage) at the same position. Rather, transcript ends vary a few nucleotides, resulting in a heterogenous population of mature transcripts (Geisberg et al. 2020; Wallace et al. 2020; Dang et al. 2024). As we performed a relatively strict analysis of TSS and TES and only kept peaks that were in at least 3 replicates, we did not assign TSS and TES to some genes because their window was wider than tolerated by our approach. Nevertheless, in most of these cases the information about the start and stop region can still be obtained by using the supplemental alignment files to assess the peak manually. We also noticed that most transcripts in *A. fumigatus* are transcribed as one isoform under normal growth conditions, with relatively limited numbers of alternative start or stop sites. How this will be affected by growth under stress conditions or during development remains an open question.

We expect such a dataset to be an engine for hypothesis generation moving forward. One exciting observation in this direction was the numerous observed examples of premature transcription termination (PTT). Although previously demonstrated in the ascomycete *S. cerevisiae*, to the best of our knowledge, this is the first report of PTT in Eurotiomycetes. We identified 42 high-confidence instances of promoter proximal PTT, but many additional examples appear to be scattered through the transcriptome for termination within the gene body. Clearly more work will be required to define both the breadth and relevance of these examples in terms of regulation. Ultimately, the definition of 5′ and 3′ ends of *A. fumigatus* transcripts will facilitate further discovery through an improved understanding of the biological underpinnings of this important pathogen. How these transcriptome features relate to other members of the *Aspergillus* species complex or fungal pathogens more broadly remains an open question that certainly deserves additional attention.

## METHODS

### Strains and culture conditions

*Aspergillus fumigatus* strain CEA10 was grown on *Aspergillus* minimal media (AMM)-agar plates for 5 days at 37°C and the spores were harvested by collection through 30-µm filters (Miltenyi Biotec). AMM was composed of 70 mM NaNO_3_, 11.2 mM KH_2_PO_4_, 7 mM KCl, 2 mM MgSO_4_, 1% [w/v] glucose, and 1 µL/mL trace element solution at pH 6.5 (1 g of FeSO_4_ • 7H_2_O, 8.8 g of ZnSO_4_ • 7H_2_O, 0.4 g of CuSO_4_ • 5H_2_O, 0.15 g of MnSO_4_ • H_2_O, 0.1 g of NaB_4_O_7_ • 10 H_2_O, 0.05 g of (NH_4_)_6_Mo_7_O_24_ • 4H_2_O, and ultra-filtrated water to 1000 mL). 50 mL of liquid AMM was inoculated with 1×10^8^ CEA10 spores and then incubated for 24 h at either 37°C or 42°C at 200 rpm. Mycelium was collected using Miracloth, washed with water, and ground with mortar and pestle in liquid nitrogen prior to downstream analyses.

### RNA-Seq and End-Seq library preparation

RNA was isolated using a TRIzol-based approach as described in (Kelani et al. 2023) with minor modifications; RNA was DNase-treated according to the manufacturer’s recommendation (ThermoFisher, RNase-free DNase) and purified using an RNA Clean & Concentrator-5 kit (Zymo Research Corporation). DNase-treated RNA was quality checked on an agarose gel and additionally the RNA Integrity Number (RIN) values were determined by Bioanalyzer (Agilent). All samples had RIN values between 9 and 9.9. To sequence 5′ and 3′ poly(A)-containing RNA ends, 3 µg of total RNA was used as input for library preparation using 5′ and 3′ RNA-End-Seq kits (Eclipse Bioinnovations) according to the manufacturer’s recommendation. In brief, for 5′ End-Seq, poly(A) RNA was enriched with oligo(dT) beads, the RNA was heat fragmented, uncapped 5′ ends were phosphorylated and digested, and the remaining 5′ terminal RNA fragments were subjected to library preparation. For 3′ End-Seq, total RNA was heat fragmented, the poly(A)-tailed fragments were isolated with oligo(dT) beads, and the AT-hybrid was digested prior to library preparation.

For poly(A)-RNA-Seq, library preparation was performed by NovoGene, UK according to standard protocols. Briefly, mRNA was purified from total RNA using poly-(T) oligo-attached magnetic beads. Following fragmentation, first strand cDNA was synthesized using random hexamer primers. The second strand cDNA was synthesized using dUTP. The directional library was considered complete after end repair, A-tailing, adapter ligation, size selection, USER enzyme digestion, amplification, and finally purification. The library was then checked by Qubit and real-time PCR for quantification and by bioanalyzer for size distribution, prior to pooling for Illumina sequencing using an Illumina Novaseq.

### Data processing and analysis

#### Total RNA-Seq

The quality of raw reads was assessed with FastQC (v0.12.0). Remaining adaptor sequences were removed with Trimgalore (v0.5.0). FastQC was applied again and the fastq files were aligned to the *A. fumigatus* CEA10 strain genome (Aspergillus_fumigatusa1163.ASM15014v1) with HiSat (v2.2.1), allowing for a maximum intron length of 5000 nt. Counts were determined by htseq-count (v0.9.1) and used as input for DeSeq2 (v2.11.40.7).

#### End-Seq

The quality of raw reads was assessed with FastQC (v0.12.0). Unique molecular identifiers (UMIs were removed with umi_tools (v0.5.5; with the setting –bc-pattern=NNNNNNNNNN for recognition of UMIs). Remaining adaptor sequences were removed with Trimgalore (v0.5.0). 5′ End-Seq reads were further treated with fastp (v0.22.0). Resulting fastq files were again checked for quality with FastQC and then aligned to the *A. fumigatus* CEA10 strain genome (Aspergillus_fumigatusa1163.ASM15014v1) with HiSat (v2.2.1), allowing for a maximum intron length of 5000 nt. The resulting bam files were converted to wig files using bam2wig (v1.6; dependent on samtools v1.14). Both read strands were then combined into a single file. To scale and transform the files into the bedgraph format, wiggletools (v1.2.11) was used with default parameters. As the 3′ end reads were antisense at this step, we swapped the strand with the “scale” option −1.

#### End-Seq Peak Calling

We performed End-Seq peak calling as previously described (Dang et al. 2024). To gain high confidence transcription start and stop sites, only peaks that were in at least 3 replicates were retained using the overlaps function of wiggletools (v1.2.11). These files were filtered to retain peaks with at least 20 reads for further analysis. For downstream processing, duplicated peaks were removed by applying bedops –merge (v2.4.41). Bedtools closest (v2.30.0) was used to assign the individual peaks to the associated genes on the chromosome. To avoid significant overlap with closely spaced genes on the opposite strand that could potentially interfere with the assignment, both the genome and the peaks were separated by strand. On the forward strand the 3′ end assignment was limited to genes upstream of the peak. Accordingly, on the reverse strand, assignment was limited to genes downstream of the peak. For the 5′ end, the gene assignment must be reverted. As such, the forward strand assignment was limited to the closest gene downstream and on the reverse strand to the closest gene upstream of the peak location. The resulting data set was then manually curated and each peak was assigned a “confidence score” of either 3 (high confidence TSS/TES), 2 (possible TSS/TES, but further experimental validation required) or 1 (unlikely a TSS/TES of a known gene in *A. fumigatus*). Genome coverage plots were made with SparK (v2.6.2) (Kurtenbach and William Harbour 2019).

### Motif analysis

For motif prediction genomic sequences were extracted from +/- 25 nt from the TSS and TES and subjected to STREME (version 5.5.7) (Bailey 2021). Randomly shuffled input sequences with maintained sequence composition were created by the STREME software itself and used as control sequences. To find binding motifs of Ssd1, PUF family proteins, and upstream translation start codons an in-house R-script requiring the following packages was used: “BiocGenerics” v0.52.0 (Biocondutor), “tibble” v3.2.1, and “tidyr” v1.3.1.

### Data and statistical analysis

Statistical analyses were performed using R version 4.4.2 (2024-10-31) in R Studio Version 2024.12.0+467 (2024.12.0+467) as described in detail in the specific methods sections or figure legends as appropriate.

## DATA ACCESS

All raw and processed sequencing data generated in this study have been submitted to the NCBI Gene Expression Omnibus (GEO; https://www.ncbi.nlm.nih.gov/geo/) with SubSeries GSE296593 and associated accession numbers GSE296590, GSE296591, and GSE296592.

## COMPETING INTEREST STATEMENT

The authors have no competing interests to declare.

## ACKNOWLEDGEMENTS

We would like to thank Amelia Barber and the members of the RNA Biology of Fungal Infections Junior Research Group for helpful discussions throughout this project. The work presented here was generously supported by the Federal Ministry of Research, Technology and Space (BMFTR: https://www.bmbf.de/), Germany, Project FKZ 01K12012 “RFIN – RNA-Biologie von Pilzinfektionen”. Additional funding support came from the Deutsche Forschungsgemeinschaft (DFG, German Research Foundation) under Germanýs Excellence Strategy – EXC 2051 – Project-ID 390713860. The funders had no role in the design, analysis, decision to publish, or preparation of the document.

## Notes

### Competing Interest Statement

The authors have declared no competing interest.

