## Supplemental Material for "Comprehensive mapping of the 5′ and 3′ untranslated regions of *Aspergillus fumigatus* reveals new insights into gene regulation"

**Running Title:** Defining *A. fumigatus* poly-(A)-enriched RNA ends.

### SUPPLEMENTAL FIGURES

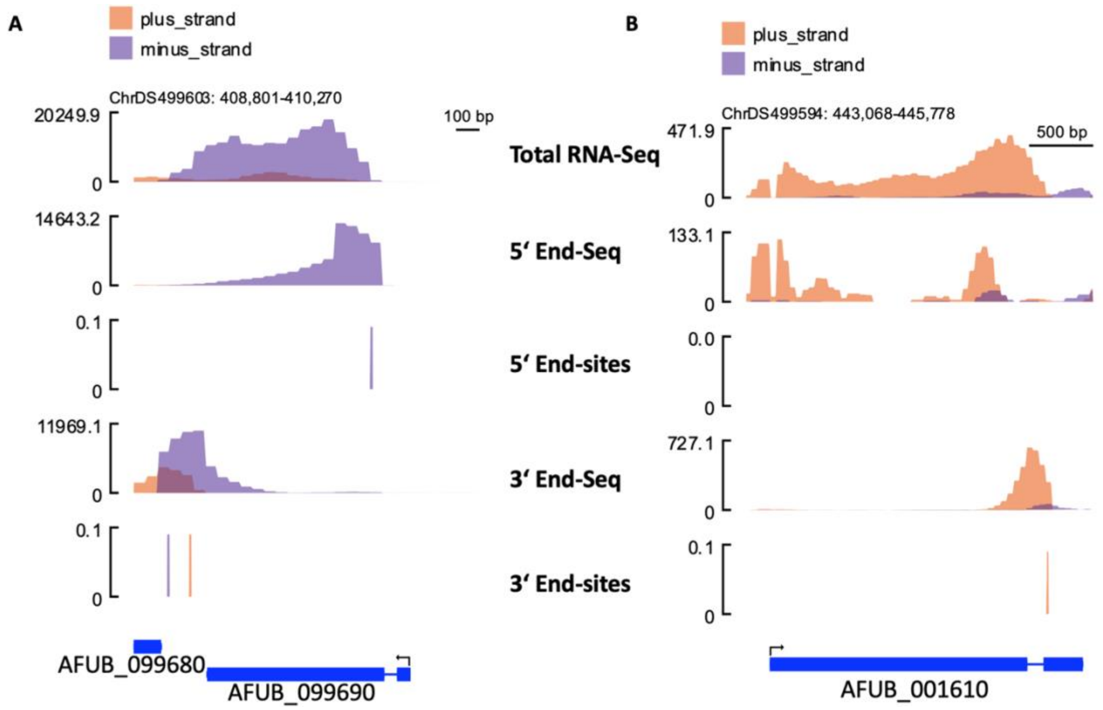


**Figure S1: UTR length bias originating from genes with predicted annotation.** (A) Read alignment of *AFUB_099680* encoding a putative transmembrane transporter and *AFUB_099690* encoding a protein of unknown function. (B) Read alignment of *AFUB_001610* encoding a protein of unknown function. In both (A) and (B), the y-axis indicates total read-counts for "-Seq" panels and arbitrary scores for "-sites". Read tracks were smoothened to improve viewability.

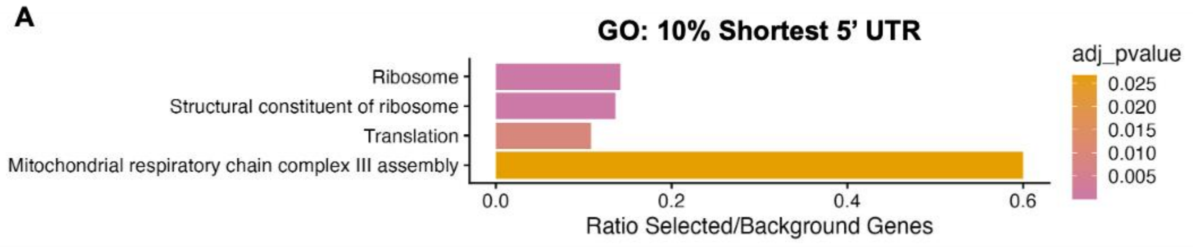
A
GO: 10% Shortest 5' UTR
Ribosome
Structural constituent of ribosome
Translation
Mitochondrial respiratory chain complex III assembly
0.0
0.2
0.4
0.6
Ratio Selected/Background Genes
adj_pvalue
0.025
0.020
0.015
0.010
0.005

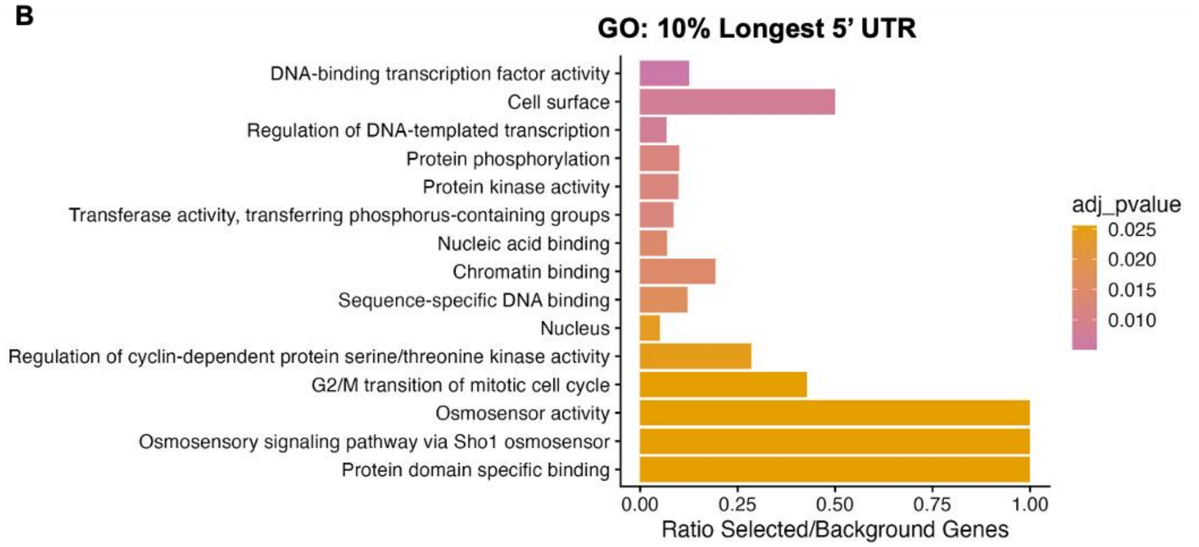
B
GO: 10% Longest 5' UTR
DNA-binding transcription factor activity
Cell surface
Regulation of DNA-templated transcription
Protein phosphorylation
Protein kinase activity
Transferase activity, transferring phosphorus-containing groups
Nucleic acid binding
Chromatin binding
Sequence-specific DNA binding
Nucleus
Regulation of cyclin-dependent protein serine/threonine kinase activity
G2/M transition of mitotic cell cycle
Osmosensor activity
Osmosensory signaling pathway via Sho1 osmosensor
Protein domain specific binding
0.00
0.25
0.50
0.75
1.00
Ratio Selected/Background Genes
adj_pvalue
0.025
0.020
0.015
0.010

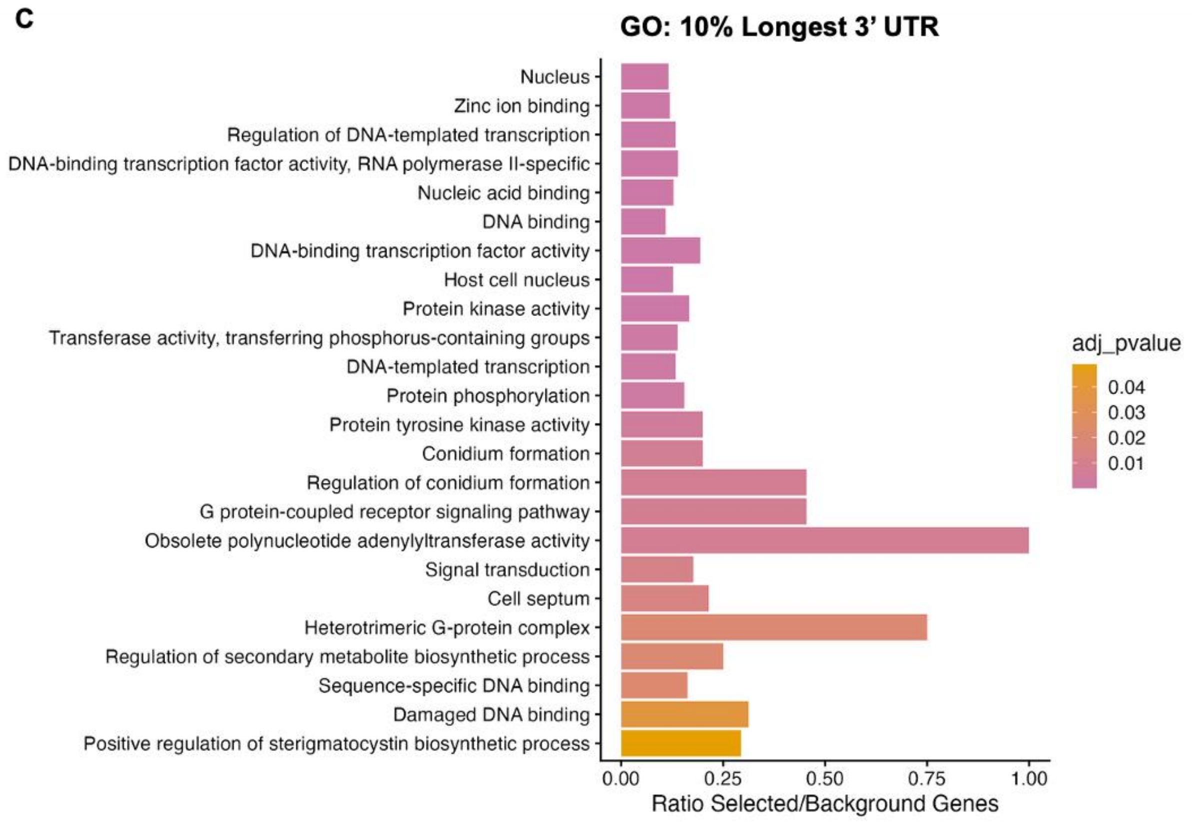
C
GO: 10% Longest 3' UTR
Nucleus
Zinc ion binding
Regulation of DNA-templated transcription
DNA-binding transcription factor activity, RNA polymerase II-specific
Nucleic acid binding
DNA binding
DNA-binding transcription factor activity
Host cell nucleus
Protein kinase activity
Transferase activity, transferring phosphorus-containing groups
DNA-templated transcription
Protein phosphorylation
Protein tyrosine kinase activity
Conidium formation
Regulation of conidium formation
G protein-coupled receptor signaling pathway
Obsolete polynucleotide adenylyltransferase activity
Signal transduction
Cell septum
Heterotrimeric G-protein complex
Regulation of secondary metabolite biosynthetic process
Sequence-specific DNA binding
Damaged DNA binding
Positive regulation of sterigmatocystin biosynthetic process
0.00
0.25
0.50
0.75
1.00
Ratio Selected/Background Genes
adj_pvalue
0.04
0.03
0.02
0.01

34

**Figure S2: GO-term analysis of the 10% shortest and longest UTRs.** Gene ontology terms associated with the genes with the (A) 10% shortest 5' UTRs, (B) 10% longest 5' UTRs, and (C) 10% longest 3' UTRs. No enrichment categories were detected for the 10% shortest 3' UTRs. GO terms were calculated with ShinyGO 0.82 (Ge et al. 2019).

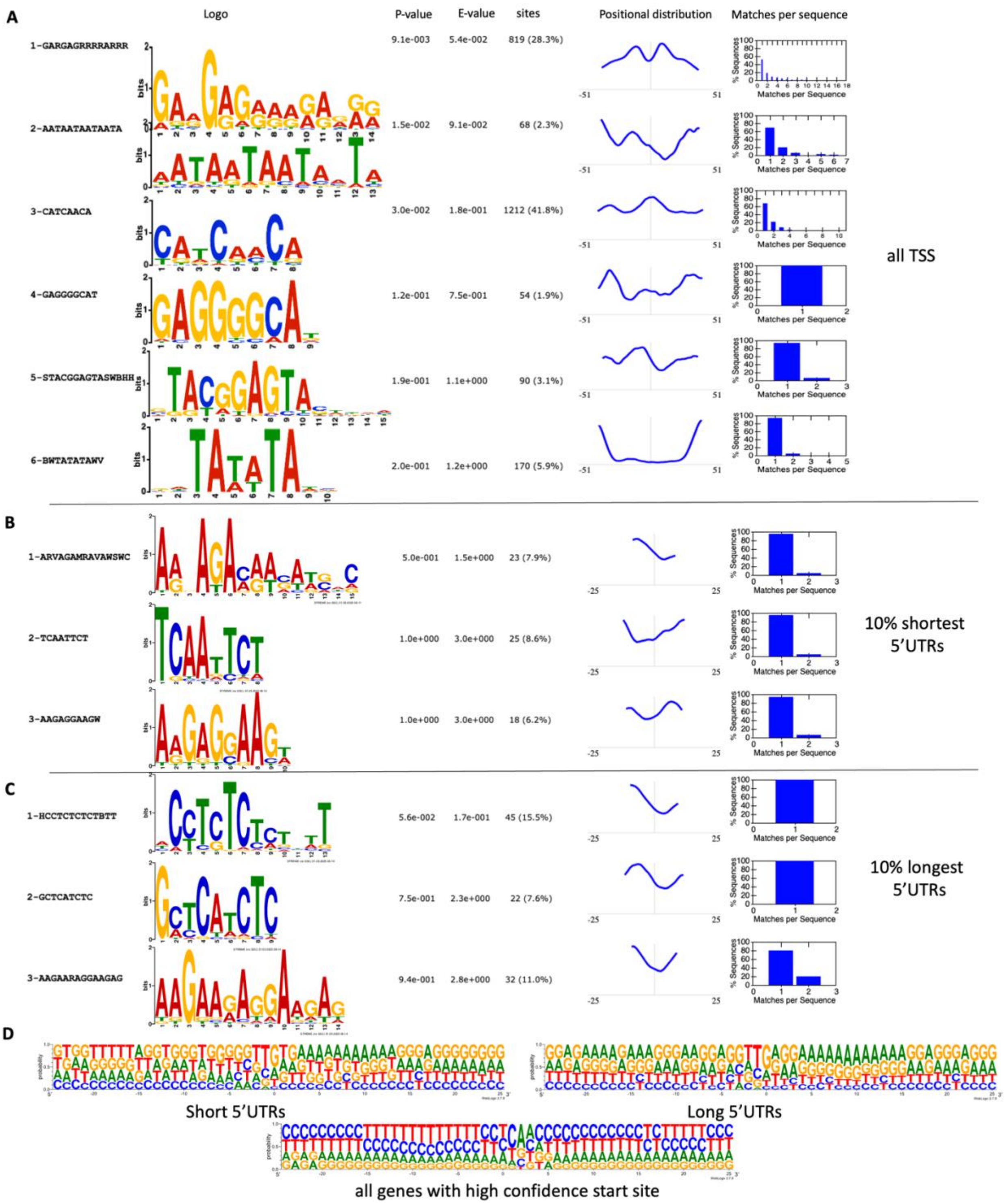


**Figure S3: Motif analysis of transcriptional start sites.** Motif prediction using STREME with genomic sequences +/- 25 nt from the transcription start sites of the (A) all 5' UTRs, (B) 10% shortest 5' UTRs and (C) 10% longest 5' UTRs. (D) WEB-logo of genomic sequences +/- 25 nt from transcription start sites for the denoted subsets or total high-confidence TES list. WEB-logo was created on https://weblogo.threeplusone.com/.

### SUPPLEMENTAL TABLES

**Table S1: DESeq2 analysis for differentially expressed genes in 42°C compared to 37°C.**

**Table S2: GO-terms of differentially expressed genes in 42°C compared to 37°C.**

**Table S3: Transcript start sites and transcript end sites for *A. fumigatus* mycelium.**

**Table S4: Number of detected uAUGs in high-confidence 5' UTRs.**

**Table S5: GO-term analysis of genes with the 10% shortest and longest 5' UTRs and 10% longest.**
